# Perturbation-aware representation learning for *in vivo* genetic screens

**DOI:** 10.1101/2025.10.15.682661

**Authors:** Florian Hugi, Tanmay Tanna, Randall J. Platt, Gunnar Rätsch

**Affiliations:** Department of Biosystems Science and Engineering, ETH Zurich, Switzerland; Department of Computer Science, ETH Zurich, Switzerland

## Abstract

CRISPR-based genetic perturbation screens paired with single-cell transcriptomic readouts (Perturb-seq) offer a powerful tool for interrogating biological systems. Yet the resulting datasets are heterogeneous—particularly *in vivo*—and currently used cell-level perturbation labels reflect only CRISPR guide RNA exposure rather than perturbation state; further, many perturbations have a minimal effect on gene expression. For perturbations that do alter the transcriptomic state of cells, intracellular guide RNA abundance exhibits a dose-response association with perturbation efficacy. We combine (i) per-perturbation, expression-only classifiers trained with non-negative negative–unlabeled (nnNU) risk to yield calibrated scores reflecting the perturbation state of single cells and (ii) a monotone guide abundance prior to yield a per-cell pseudo-posterior that supports both assignment of perturbation probability and selection of affected gene features. To obtain a low-dimensional representation that allows for the accurate reconstruction of gene-level marginals for counterfactual decoding, we train an autoencoder with a quantile–hurdle reconstruction loss and feature-weighted emphasis on perturbation-affected genes. The result is a perturbation-aware latent embedding amenable to downstream trajectory modeling (e.g., optimal transport or flow matching) and a principled probability of perturbation for each non-control cell derived jointly from its guide counts and transcriptome.

## 1 Introduction

CRISPR–Cas9–mediated genetic perturbation screens have transformed the study of cellular systems, and their coupling to single-cell state readouts enables detailed understanding of gene function^1^ and unlocks direct modeling of the consequences of genetic modulation. Although most computational methods have been developed around large-scale *in vitro* datasets^2^, recent experimental methods provide rich transcriptomic measurements from *in vivo* perturbation screens^3^ — datasets that incorporate cellular and environmental context, capture heterogeneity, and carry immediate relevance for health (Figure 1A). However, *in vivo* single-cell CRISPR screens differ materially from their *in vitro* counterparts: cell cohort sizes are modest, cellular compositions are heterogeneous, and guide RNA labels are an indicator only of exposure and not perturbation efficacy. In this regime, while control-labeled cells generally constitute trustworthy negatives, guide-labeled cells form a mixture of non-perturbed, control-like cells and true positives, since editing efficiency and outcomes are heterogeneous. Furthermore, even when editing occurs, some perturbations are contextually inert, producing no detectable transcriptomic shift and leaving edited cells indistinguishable from controls.

**Figure 1:**
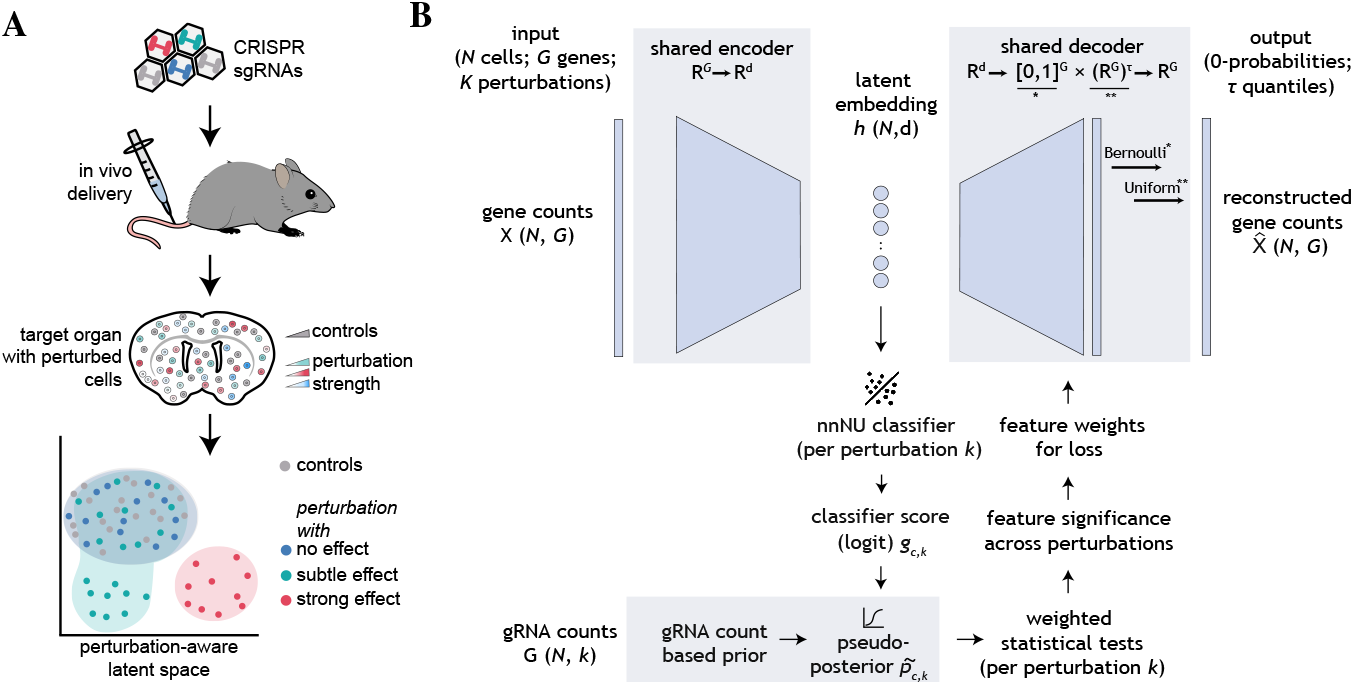
Overview of biological setting and method. **A**. *in vivo* CRISPR guide RNA delivery yields control and guide-labeled cells with variable perturbation efficacy; aim: a perturbation-aware latent representation. **B**. Model architecture.

The NeurIPS 2025 Workshop on AI Virtual Cells and Instruments: A New Era in Drug Discovery and Development (AI4D3 2025), San Diego, California, USA, 2025.

Recent studies have shown that for perturbations that do induce a transcriptomic shift, the number of intracellular guide molecules can serve as a proxy for efficacy, i.e., there is a guide dose-dependent perturbation response^4^; however, this association is noisy and guide abundance is better treated as a prior signal rather than a hard label.

Our objectives are twofold. First, we aim at constructing perturbation-aware latent representations suitable for trajectory modeling that support the reconstruction of realistic gene marginals. Second, we want to assign each non-control cell a probability of perturbation by integrating transcriptomic evidence with guide abundance. We realize these goals within a single iterative loop that generates refined embeddings, feature reconstructions, and per-cell perturbation probabilities (Figure 1B).

This loop involves three interacting pieces: (a) an autoencoder (AE) trained using a quantile–hurdle loss, (b) per-perturbation classifiers trained with a non-negative negative–unlabeled (nnNU) risk (an adaptation of non-negative positive unlabeled risk^5^) to handle asymmetric label noise and class imbalance, and (c) a monotone guide prior derived from guide abundance. During each iteration, we up-weight genes whose signal is salient for perturbation-induced variance and fit the autoencoder, encode cells, train the per-perturbation classifiers in latent space, and fuse classifier logits with the guide prior in logit space to obtain per-cell pseudo-posteriors. These posteriors drive weighted significance testing to update gene priorities, which in turn refresh the reconstruction weights for the next round. This iterative scheme jointly refines embeddings and pseudo-posteriors, progressively concentrates model capacity on perturbation-responsive features, and yields principled per-cell probabilities along with a perturbation-aware representation suitable for trajectory modeling.

## 2 Dataset

Here, we focus on a publicly available *in vivo* Perturb-seq dataset profiling how mouse cortical neurons respond to perturbations in genes associated with DiGeorge syndrome^3^. The screen targets 29 genes, with two guide RNAs per gene, and reports guide abundance and transcriptomes at a singlecell resolution. Three major neuronal classes are assayed with sufficient cell numbers — interneurons, deep-layer neurons, and superficial-layer neurons, with approximately 200 cells per perturbation per cell type (Table 1). Four perturbations exhibit statistically significant transcriptomic effects based on the reporting criteria of the original study. Control-labeled cells contain non-targeting guides and serve as reliable negatives; guide-labeled cells for a perturbation are heterogeneous mixtures that include both perturbed and non-perturbed states, and are used accordingly in our negative-unlabeled training setup and downstream analyses.

**Table 1:**
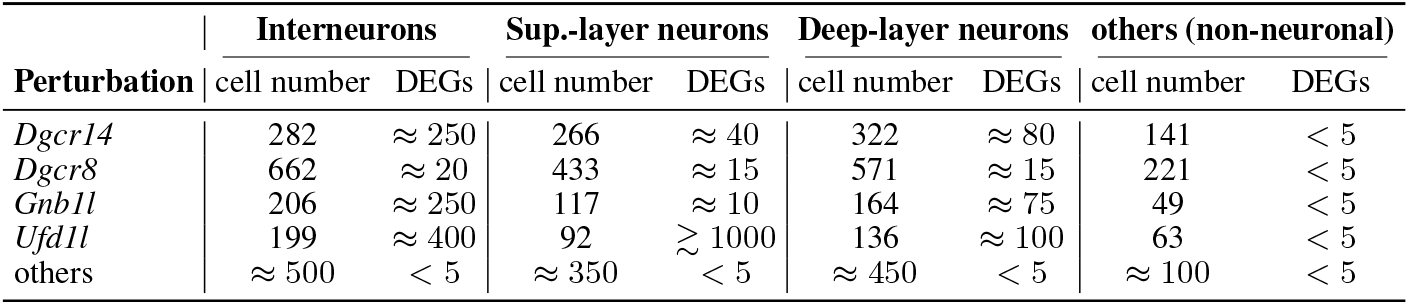
Cell counts and differentially expressed gene (DEG) numbers reported in *Santinha et al*.^3^.

## 3 Problem setting

Let *N* cells be indexed by *c* ∈ {1, …, *N*}, *G* genes by *j* ∈ {1, …, *G*}, and *K* perturbations by *k* ∈ {1, …, *K*}. Each cell has a log-normalized expression vector *x*_*c*_ ∈ ℝ^*G*^. For perturbation *k*, the cell carries guide counts 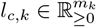 across *m*_*k*_ guides (zero for controls). Let *d*_*c,k*_ ∈ {0, 1} indicate whether any guide for *k* is observed, and let *y*_*c,k*_ ∈ {0, 1} denote the *unobserved* true perturbation state. We assume

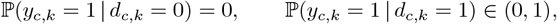

i.e., labeled controls are reliable negatives, while guide-labeled cells are a mixture of perturbed and unperturbed states. Our goal is to estimate for each (*c, k*) a probability 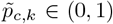 that the cell is perturbed by *k*, and to learn a map *h*: ℝ^*G*^ → ℝ^*d*^ and decoder Dec: ℝ^*d*^ → ℝ^*G*^ that allow the accurate reconstruction of log-normalized gene expression counts, with ℝ^*d*^ being a low-dimensional latent space which captures perturbation structure and enables trajectory modeling from control to perturbed states.

## 4 Methods

### 4.1 Latent embedding with quantile–hurdle decoding

Conventional AE/VAE formulations trained with mean-squared-error (MSE) objectives match only conditional means^6^ and therefore fail to reproduce the zero-inflated, heavy-tailed gene-count marginals characteristic of single-cell data. Other methods (e.g., scVI^7^) use parametric models which operate on raw counts under a prespecified likelihood (negative binomial/Poisson) that fixes the mean–variance relationship and tail behavior, an assumption that can be misspecified for heterogeneous *in vivo* datasets and is thus restrictive for marginal reconstruction. Here, we train an encoder *h*: ℝ^*G*^ → ℝ^*d*^ to produce low-dimensional embeddings *z*_*c*_ = *h*(*x*_*c*_) and a decoder that, for each gene *j*, predicts (i) a zero-event probability 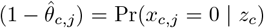 to capture the zero-inflated nature of single-cell RNA counts and (ii) a set of conditional quantiles 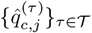 of *y*_*c,j*_ = log(1 + *x*_*c,j*_) given *x*_*c,j*_ > 0. This enables faithful reconstruction of log-normalized single-cell expression beyond feature means, with realistic, zero-inflated gene marginals, without imposing explicit parametric assumptions on the count distribution. Perturbation-affected features are emphasized through an iterative learning process which is described in subsequent sections that couples autoencoder training with perturbation-guided feature weighting.

#### Hurdle component

For each gene *j*, let *b*_*c,j*_ = **1**{*x*_*c,j*_ > 0} and let 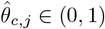 be the decoder’s estimate of ℙ (*x*_*c,j*_ > 0 | *z*_*c*_). The zero-inflation term is

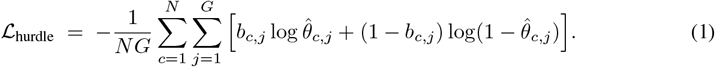

#### Quantile component

Conditioned on *x*_*c,j*_ > 0, we fit a set of conditional quantiles of a transformed magnitude *y*_*c,j*_ = log(1 + *x*_*c,j*_). For a finite set 𝒯 ⊂ (0, 1) of quantile levels, the set 𝒫 = {(*c, j*) | *x*_*c,j*_ > 0}, and decoder predictions 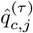, the quantile loss is

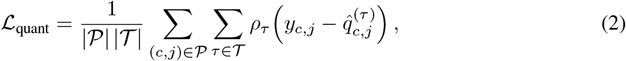

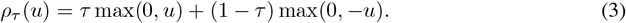

#### Feature-weighted training

To focus model capacity on selected features, we introduce nonnegative per-gene weights 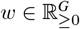. Base importances 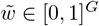 (from any source; here iteratively updated) are rescaled as 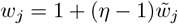 with *η* > 1. Let 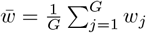 denote the mean weight. For either loss component • ∈ {hurdle, quant}, we apply weighting as

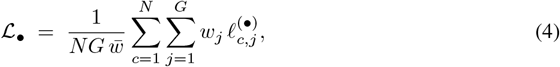

where 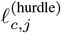 is the Bernoulli (zero) term and 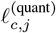 is the quantile term (averaged over *τ* ∈ 𝒯). Increasing *w*_*j*_ prioritizes accurate reconstruction of feature *j*, which in turn encourages the encoder to allocate latent capacity to the variation captured by that feature.

#### Full objective

The overall training objective combines the quantile and hurdle components with an *ℓ*_2_ penalty ℒ_reg_ on the latent representations *z*_*c*_; *λ*_1_, *λ*_2_ ≥ 0 are loss weights.

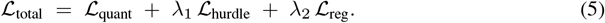

#### Feature reconstruction

To generate reconstructed features 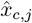 from the decoder outputs, we combine the hurdle and quantile components. First, a Bernoulli draw with success probability 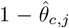 determines whether the feature is set to zero. Conditioned on being nonzero, we draw *u* ∼ Uniform(0, 1) and map it to a value between adjacent predicted quantiles 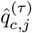 using piecewiselinear interpolation.

### 4.2 Cell-state-based perturbation classification under negative–unlabeled regime

As discussed before, we work in a negative–unlabeled (NU) regime, broadly similar to a positive– unlabeled (PU) regime^5^: controls are reliable negatives, while guide-labeled cells mix perturbed and unperturbed states. For each perturbation *k*, we train a parametric *logit* score function *f*_*k*_: ℝ^*d*^ → ℝ implemented as a lightweight MLP with a single hidden layer, which outputs a score *g*_*c,k*_ = *f*_*k*_(*z*_*c*_) that represents expression-based perturbation efficacy. A larger *g*_*c,k*_ indicates stronger evidence that *y*_*c,k*_ = 1 based on the latent cell state *z*_*c*_. Training uses control cells as a set of reliable negatives 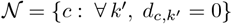 and an *unlabeled* pool 𝒰_*k*_ = {*c*: *d*_*c,k*_ = 1}, which mixes positives and negatives; *d*_*c,k*_ ∈ {0, 1} is an indicator for whether guide *k* is observed in cell *c*.

#### nnNU risk

We define the positive and negative logit losses as *ℓ*^+^(*g*) = softplus(−*g*) and *ℓ*^−^(*g*) = softplus(*g*). Let 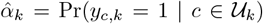 denote the estimated positive fraction within the unlabeled pool. We then minimize the non–negative NU empirical risk

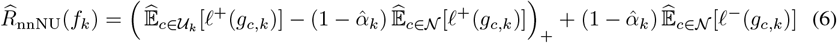

where (·)_+_ prevents the negative risk from becoming spuriously small in the presence of mislabeled positives. Because 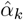 is unknown, we initialize it at a conservative upper bound and iteratively refit it until the mean score on reliable negatives—after a logistic transform—falls below a small target.

#### Monotone guide-based prior

We treat guide abundance as monotonic evidence for perturbation efficacy. For each perturbation *k*, per-cell guide counts are aggregated and mapped via a rank-to-*z* transform to yield *u*_*c,k*_ (control cells are mapped to a fixed lower bound, e.g., −5, and guide-labeled cells are mean-centered at 0 for cross-perturbation comparability). A single nonnegative scale parameter *τ*_*u*_ ≥ 0 converts this signal into a log-odds increment, guide logit contribution: *τ*_*u*_ *u*_*c,k*_. (7)

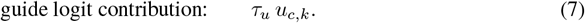

#### Logit-space fusion to a pseudo-posterior

We combine the expression-based score *g*_*c,k*_ with the guide-based logit contribution (Equation 7) additively in log-odds space and map the result through the logistic function to obtain a pseudo-posterior (i.e., not a full Bayesian posterior but a calibrated heuristic) in [0, 1]:

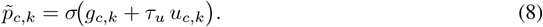

The mapping is monotone in each argument; when 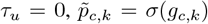; for ambiguous cells (*g*_*c,k*_ ≈ 0), the guide signal *u*_*c,k*_ materially shifts the probability; and for confident cells (large |*g*_*c,k*_|), sigmoid saturation attenuates the incremental influence of the prior.

#### Training procedure

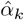 is unknown and can range from 0 to 1, reflecting both the perturbed fraction among unlabeled cells and whether the perturbation effect is detectable in latent space. To estimate 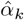, we start from a conservative upper bound and iteratively decrease it, refitting until the mean classifier score *g*_*c,k*_ on reliable negatives in a held-out validation split falls below a small target; negatives thus serve as calibration controls.

### 4.3 Weighted significance testing and feature prioritization

We quantify perturbation-driven effects on gene expression by replacing hard guide-based labels (which are noisy) with per-cell pseudo-posteriors 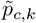 and conducting weighted tests against controls. For each perturbation *k*, we perform per-gene weighted Welch tests that compare a pseudoposterior–weighted test group to controls.

Let 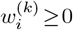 denote per-cell frequency weights for the test group of perturbation *k*, here derived from the pseudo-posterior as 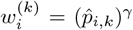 (with *γ* > 0). Controls from the same cell type form the reference group and are used unweighted (weight = 1). Weighted means and variances for the test group are computed using 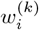, and the Kish effective sample size 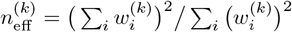 is used in the standard error and in the Welch–Satterthwaite degrees of freedom. This yields two-sided *p*-values 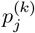 for each gene *j*. For each *k*, we apply a Bonferroni correction over the *m* tested genes. Finally, adjusted *p*-values are mapped by a fixed non-increasing transform *ϕ*: [0, 1] → [0, 1] to priorities 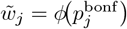, which serve as base importances for the next iteration of feature-weighted training. In this work, for the sake of simplicity, we use the hard threshold *ϕ*(*u*) = **1**{*u* ≤ 0.05}.

## 5 Overall training procedure

We estimate autoencoder parameters and per-cell pseudo-posteriors in a coupled, iterative loop; the full algorithm is provided in the appendix.

### Initialization and warm start

Since no pseudo-posterior is available at initialization, we first utilize the guide-based priors to perform weighted feature testing (Subsection 4.3) and obtain an initial set of perturbation-aware genes. We train the autoencoder with the feature-weighted quantile–hurdle objective (Equation 5) with these genes up-weighted. On the resulting embeddings, we train per-perturbation classifiers and compute pseudo-posteriors via logit fusion with the guide-based prior (Equation 8), keeping *τ*_*u*_ > 0 to stabilize ambiguous cells and to provide a reliable initial ranking.

### Iterative refinement and convergence

In subsequent iterations, we repeat the following cycle: refit the autoencoder with updated feature weights, re-encode cells, retrain per-perturbation classifiers in latent space, recompute pseudo-posteriors, and update gene priorities via weighted significance testing. We restrict attention to perturbations that are not inert (screened by a fixed top-half posterior gap relative to controls). Across iterations, we monotonically decrease the guide-prior scale (*τ*_*u*_ ↓ 0) while increasing the feature-emphasis scale (*η*) up to a prespecified cap, thereby shifting reliance from prior-driven warm starts to data-driven evidence. By design, the loop runs for a fixed number of iterations *T*_max_ while ramping *η* to *η*_max_ and annealing *τ*_*u*_ to *τ*_min_; we then return the trained autoencoder, per-perturbation classifiers, final pseudo-posteriors 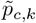, and the selected gene set with final weights—outputs that can be used for deconstructions and downstream trajectory modeling.

## 6 Results

### 6.1 Quantile-hurdle decoding

Our quantile–hurdle objective preserves empirical gene marginals—including zero inflation and tail behavior—upon decoding, outperforming standard AE/VAE methods trained with mean-squarederror objectives, without utilizing parametric count likelihoods (NB/Poisson) that typically require raw, unnormalized counts and impose fixed mean–variance relationships. To illustrate, we present Q–Q plots and empirical marginals for three exemplar genes, comparing the original distributions against reconstructions from (i) an AE trained with mean squared error and (ii) our AE with the quantile–hurdle loss (Figure 2). The example genes are representative of three different classes: a high-expression gene with negligible zero inflation, a low-expression gene with pronounced zero inflation, and a median-expression gene with moderate zero inflation.

**Figure 2:**
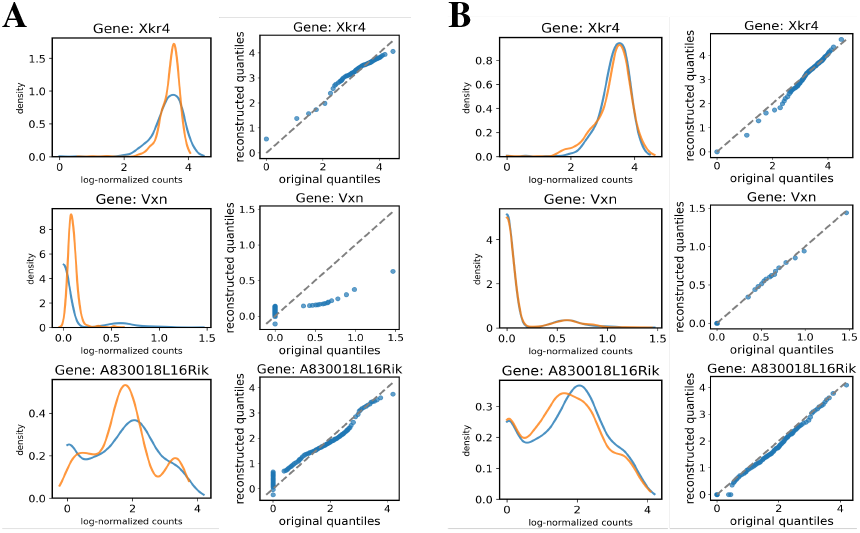
Feature marginals and Q-Q graphs showing original and reconstructed log-transformed normalized feature counts of three highly variable genes for **A**. AE trained with mean squared error loss **B**. AE trained with quantile-hurdle loss.

### 6.2 nnNU classification

When perturbation-driven variance is adequately captured in the latent codes, training per-perturbation *logit* score functions using nnNU risk and fusing them with guide-count priors *σ*(*u*_*c,k*_) yields pseudo-posteriors that distinguish truly perturbed cells from guide-labeled but unperturbed cells. We illustrate this with *Ufd1l*, a perturbation that induces a strong transcriptional shift (Table 1). In latent space the perturbation is well separated, yet a subset of guide-labeled cells clearly overlaps with the control distribution (Figure 3A). The guide-based prior *σ*(*u*_*c,k*_)—a function of guide RNA abundance—improves separation (Figure 3B), confirming recent studies^4^. The per-cell pseudoposterior, obtained by combining the classifier score with the guide-based prior, further refines the boundary and provides a high-confidence population of truly perturbed cells (Figure 3C).

**Figure 3:**
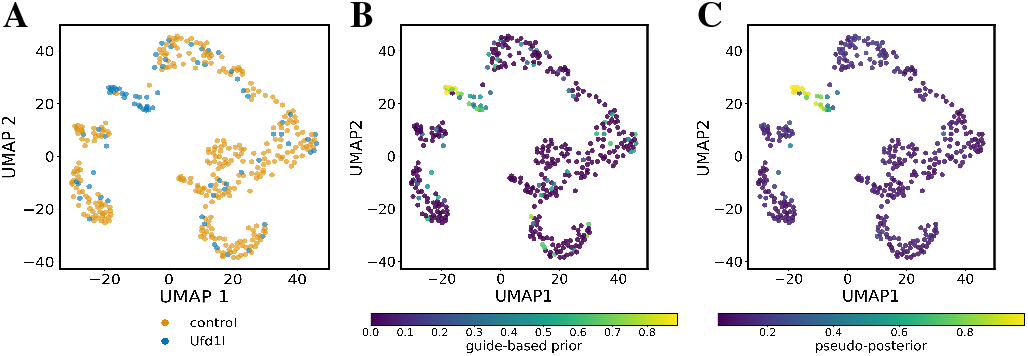
UMAP projection of latent codes of control and *Ufd1l*-perturbed cells from a validation set held out during training colored by **A**. guide label **B**. guide-based prior **C**. pseudo-posterior.

### 6.3 Iterative training

We evaluate the iterative refinement procedure (Section 5) on the interneuron subset of the dataset (Section 2)—cells transduced with either an inert control guide RNA or any of the 29 tested perturbations—and benchmark against a single, unweighted autoencoder trained on all features with a downstream classifier on its latent space. In the baseline embeddings from an unweighted quantile–hurdle AE, only *Ufd1l* forms a clearly separated cluster, whereas the remaining perturbations are largely intermixed with controls (Figure 4A). Using our approach, the embeddings become progressively more structured and significant perturbations resolve into distinct regions of latent space (Figure 4B).

**Figure 4:**
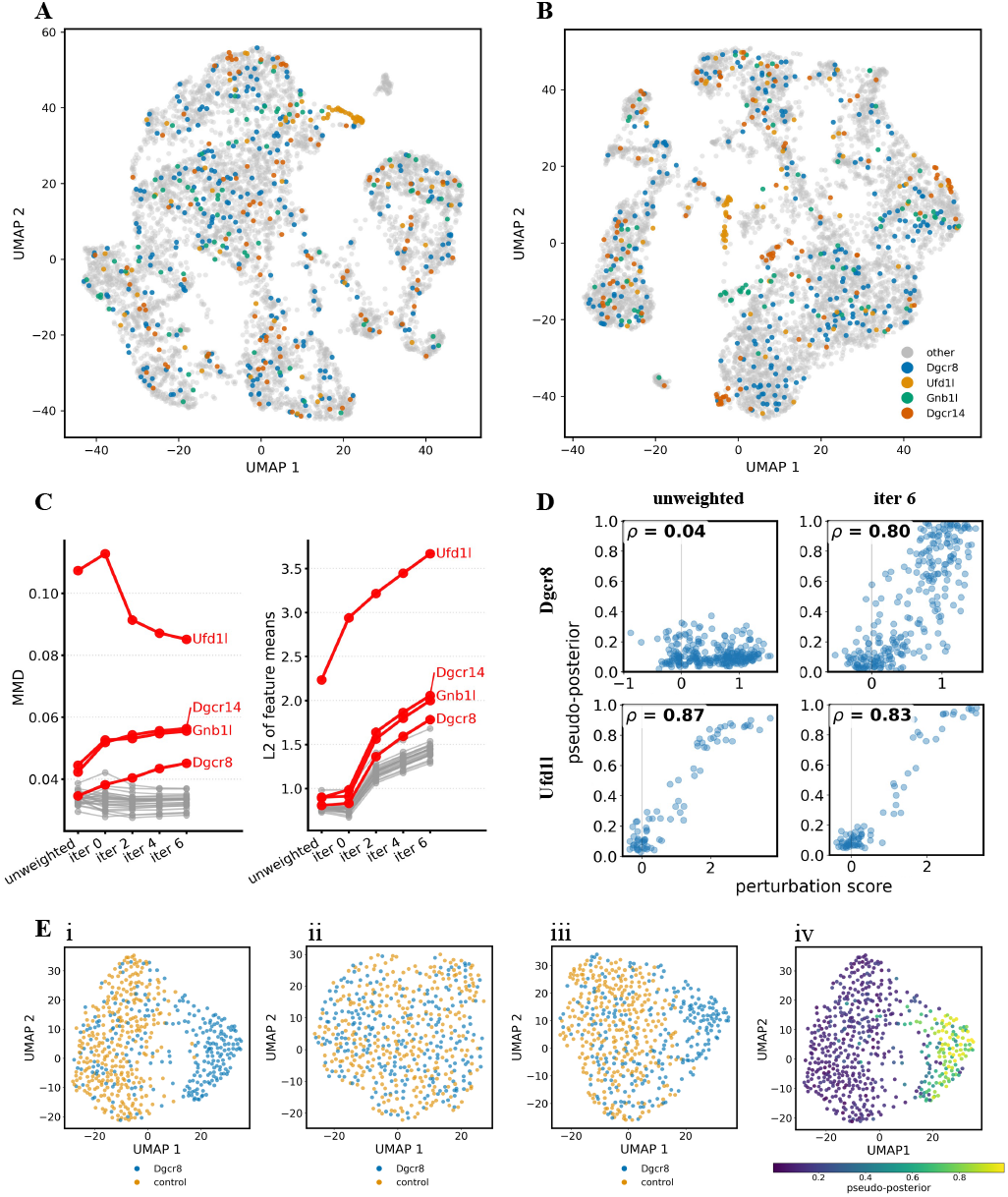
Iterative refinement of perturbation–aware embeddings in interneurons. **A**. UMAP of unweighted quantile–hurdle AE embeddings; only *Ufd1l* separates cleanly from controls. **B**. UMAP after iterative training with feature weighting. **C**. Quantification across iterations: kernel MMD (left) and *L*_2_ distance between latent means (right) vs. controls increase for effective perturbations (*Ufd1l, Dgcr14, Gnb1l, Dgcr8*). **D**. Association between pseudo–posterior 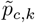 and a marker–based perturbation score strengthens from baseline to final iteration (iter 6) (Spearman *ρ* shown) for *Dgcr8* and is retained for *Ufd1l* **E**. Marker–space reconstructions for *Dgcr8*: (i) original marker UMAP; (ii) unweighted AE reconstructions (separation lost); (iii) iterative method reconstructions (separation recovered); (iv) original cells recolored by 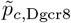.

We quantify separation in the latent space using two complementary metrics: the kernel maximum mean discrepancy (MMD) with a Gaussian radial kernel and the *L*_2_ distance between latent-space feature means for each perturbation versus controls (Figure 4C). In the baseline setting, separation is evident only for *Ufd1l*; after multiple iterations, perturbations with subtler effects, such as *Dgcr8*, are also identified. To relate latent separation to expression-level signal, we define a perturbation score for each cell by *z*-scoring the top 25 markers per perturbation from Santinha *et al*.^3^ and aggregating across markers. The association between the pseudo-posterior 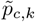 and this perturbation score strengthens under our procedure—*Ufd1l* exhibits strong baseline concordance, whereas for *Dgcr8*, this becomes pronounced only after iterative refinement, with corresponding sharpening of 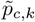 (Figure 4D). Finally, reconstructed features retain perturbation-specific signatures under the proposed training (Figure 4Ei/iii), as illustrated for *Dgcr8*, the weakest effective perturbation in this dataset, while the unweighted AE baseline fails to recover these effects (Figure 4Eii).

## 7 Discussion

There has been substantial progress in modeling cellular responses to genetic perturbations: some approaches have focused on specifying parametric functions for predicting perturbed states^8–10^ while, more recently, empirically driven “foundation”–style models that learn broadly transferable response mappings from large-scale multi-cohort datasets have gained prominence^11–13^, with most of these efforts being applied to *in vitro* data. The *in vivo* setting introduces greater heterogeneity and label uncertainty, requiring methods that are statistically robust and biologically grounded.

Our contribution addresses this regime by constructing perturbation–aware latent representations while exploiting the asymmetry of labels intrinsic to *in vivo* Perturb–seq: control cells comprise a reliable negative class, and per–perturbation *logit* score functions trained using nnNU risk address mislabeling in the perturbed class. We also leverage a monotone guide–based prior as orthogonal evidence reflecting molecular dosage, and logit–space fusion—with a prevalence correction—yields interpretable pseudo–posteriors 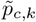 for downstream testing. On the representation side, a quantile–hurdle decoder prioritizes reconstruction of gene marginals (zero inflation and tail behavior), which is essential when projecting latent manipulations back to expression space; feature weighting *w*_*j*_ concentrates capacity on perturbation–responsive genes, enhancing the resolution of perturbation structure in latent space.

### Assumptions, limitations, and alternatives

Our approach assumes (i) that control-labeled cells constitute reliable negatives and that non-perturbed, guide-labeled cells are contained within the control distribution, (ii) a stable mapping from guide abundance to efficacy that justifies a monotone prior. Practical limitations include sensitivity to the initialization of the positive fraction 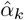 (we use a conservative upper bound 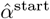) and to the schedule for annealing the guide prior; and the absence of end–to–end uncertainty propagation from 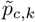 into the embedding. Further, calibration across perturbations can drift when classifiers are trained independently; common–scale calibration can address this but is not yet incorporated into our present analysis. Two methodological alternatives can be pursued: an EM–based loop that treats 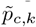 as latent posteriors rather than pseudo–posteriors, and a SCAR–based formulation in place of nnNU to remove the need to estimate the positive fraction 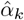. Bayesian hierarchical variants could further regularize per–perturbation scales and propagate uncertainty.

### Future directions

Our refined latent space is amenable to downstream trajectory modeling of perturbation effects using approaches such as optimal transport or flow matching, with the quantile–hurdle decoder providing a calibrated map back to gene space. Extending the framework to multitask calibration across *k*, integrating stronger priors on guide efficacy, and coupling uncertainty–aware decoding with soft responsibilities are promising next steps.

### Broader impact statement

This work advances computational methodology for small, heterogeneous *in vivo* single-cell CRISPR screens by integrating molecular priors with statistically robust classification and reconstruction objectives in a biologically principled manner. The approaches are designed to improve downstream inference tasks without increasing experimental burden and may thus facilitate safer, more informative *in vivo* studies.

## 9 Appendix

### 9.1 Data processing

We downloaded the pre-processed single-cell CRISPR dataset described in Santinha et al.^3^ from scPerturb.^14^ Cells with fewer than 6,000 total counts or fewer than 2,700 detected genes were removed; these thresholds were chosen from empirical count/feature distributions. Genes detected in fewer than 3 cells were excluded, as were mitochondrial genes. To filter uniformly uninformative features, we computed the mean expression for each gene within every experimental condition (i.e., control and each perturbation were considered separately) and took the maximum of these means across conditions; genes whose maximum mean was less than 0.4 were discarded. This removes genes that are essentially not expressed in any condition but keeps features that are expressed even in small groups of perturbation-relevant cells. Libraries were then normalized by total counts to 10,000 per cell and log1p transformed.

### 9.2 Algorithm

#### Algorithm 1

Iterative weighted quantile–hurdle AE with nnNU classification

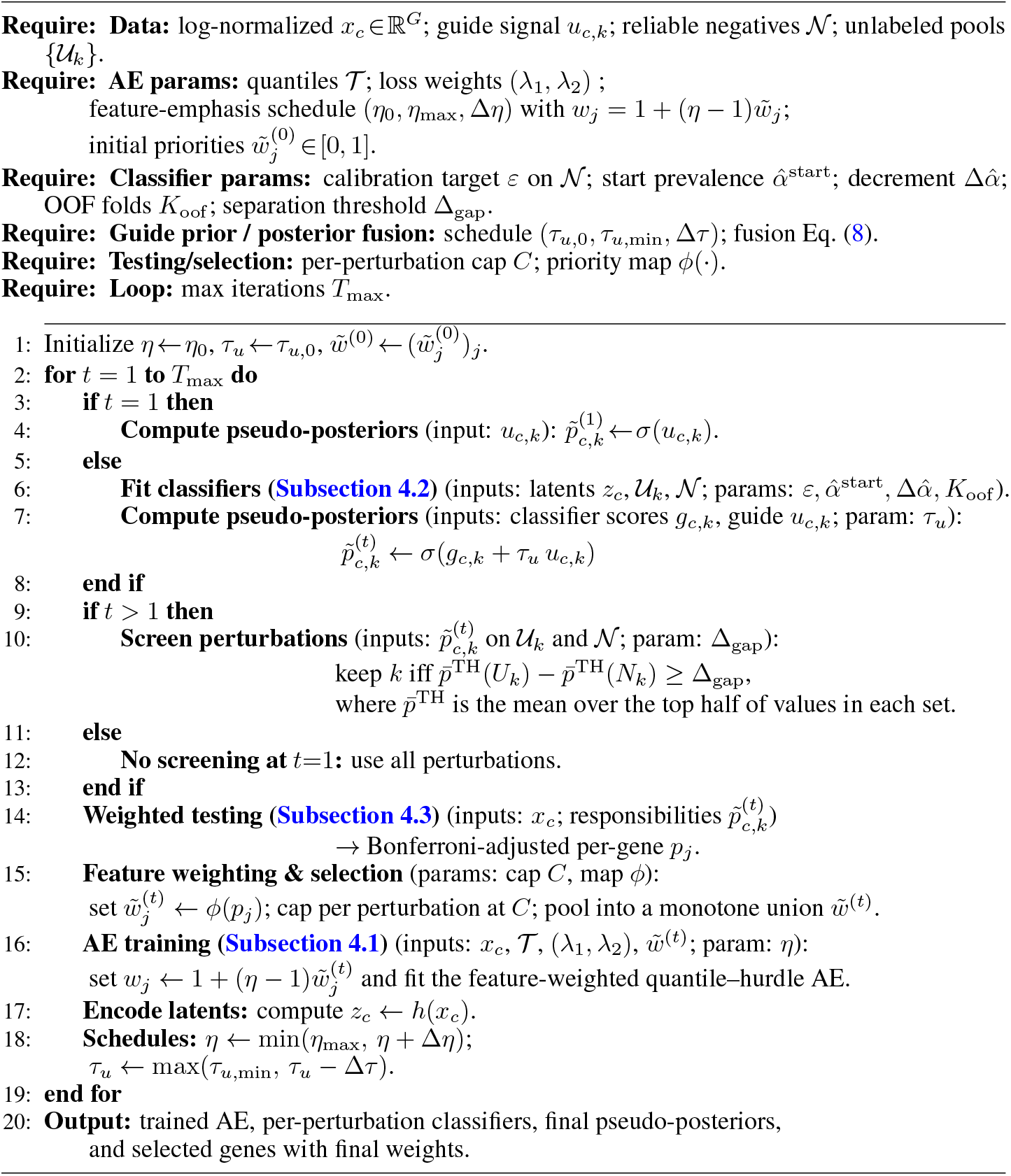

### 9.3 Evaluation metrics

We quantify separation in latent space using two complementary metrics computed on the *validation* split: kernel maximum mean discrepancy (MMD) with a Gaussian radial kernel and the *ℓ*_2_ distance between latent-space feature means (control vs. each perturbation). Let *X* = {*x*_*i*_} denote validation latent codes for control cells and *Y* = *y*_*j*_ those for a given perturbation. For each perturbation, we enforce equal sample sizes by drawing *n* = {min |*X*|, |*Y*|, min_*p*≠control_ |*Y*_*p*_|} cells from *X* and from the target *Y*; sampling is without replacement when possible and with replacement otherwise. We repeat this procedure *R* times (default *R* = 100) and report the average metric across repetitions.

#### Kernel MMD (Gaussian RBF)

For each repetition we compute the (biased) MMD^2^ estimate

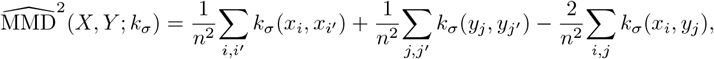

with 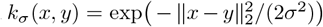. Bandwidths are chosen data-adaptively from controls: we compute the median pairwise Euclidean distance (on up to 500 randomly selected validation controls), set a base scale 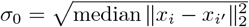, and evaluate a small kernel bank 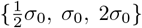. The reported MMD for a perturbation is the mean of 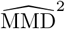 across these bandwidths and across the *R* repetitions.

#### *ℓ*_2_ distance of feature means

Within each repetition we compute

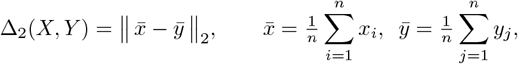

i.e., the Euclidean norm of the difference between control and perturbation mean vectors in latent space. The final *ℓ*_2_ score is the average of Δ_2_ over the *R* repetitions.

For completeness, the summary table additionally records, per perturbation, the validation set size used (*n*_val, pert_) and the target draw size *n* for the equal-size comparisons.

### 9.4 Training details

The autoencoder (AE) was trained with two-layer MLP encoder/decoder (hidden units [512, 512]) and latent dimension *d* = 50. The reconstruction objective followed Eq. (5) with hurdle (crossentropy) weight *λ*_1_ = 0.5 and a small latent *ℓ*_2_ penalty *λ*_2_ = 10^−5^. The decoder predicted | 𝒯 | = 44 conditional quantiles (from 0.001 to 0.999), and reconstructions used the sample-based procedure described in the Methods. Minibatches were of size 256, and the AE was optimized with Adam (learning rate 10^−3^). As a warm start, we trained the AE for 500 iterations using prior-based feature weights from the initialization round (Sec. 4.3); after each outer iteration we continued AE training for another 500 iterations on the updated weights. Feature emphasis used the weight map 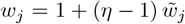; we set *η*^(1)^ = 10^2^ at warm start and then increased it by a factor of 10 until reaching the cap *η*_max_ = 10^4^ where it was held fixed thereafter.

Expression-only probe classifiers *f*_*k*_ (one per perturbation) operated on fixed latents and consisted of a single hidden layer with 16 units and dropout 0.5. They were trained for 7000 optimization steps with AdamW (learning rate 10^−4^, weight decay 5 × 10^−4^) under the nnNU risk (Eq. (6)); the best-validation checkpoint was used for predictions. For each cell-type–perturbation pair, the positive prevalence was initialized at 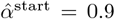 and decreased in steps of 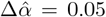 until the mean pseudo-posterior on validation controls satisfied the calibration target ≤ 0.15; the resulting 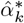 was then held fixed to compute *K* = 4 out-of-fold (OOF) predictions on the training split. Progression required an OOF top-half mean gap Δ_gap_ ≥ 0.15 between guide-exposed and control cells.

Guide information entered only via the logit-space fusion in Eq. (8). We set the guide scale to 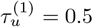 in the first outer iteration to stabilize early ranking and ablated 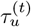 in each subsequent iteration (*t* ≥ 2), i.e., pseudo-posteriors 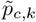 were expression-derived after the latent space had been refined.

Weighted Welch tests (Sec. 4.3) used responsibilities 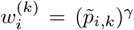 with *γ* = 1.0. We applied Bonferroni correction at *α* = 0.05 for both the initialization (prior-based) pass and the OOF-based pass, and capped the number of up-weighted genes at 100 per (cell-type *×* perturbation) category. Categories with at least 10 significant genes were oversampled by a factor of 10 during AE training. The outer loop ran for up to 6 iterations.

